# Interaction range of common goods shapes Black Queen dynamics beyond the cheater-cooperator narrative

**DOI:** 10.1101/2024.07.16.603646

**Authors:** Matthew S. Fullmer, Bram van Dijk, Nobuto Takeuchi

**Affiliations:** University of Auckland, Biological Sciences, New Zealand; Utrecht University (Theoretical Biology), the Netherlands

## Abstract

Dependencies among microorganisms often appear mutualistic in the lab, as microbes grow faster together than alone. However, according to the Black Queen (BQ) hypothesis, these dependencies are underpinned by the evolutionary benefits from loss-of-function mutations when others in the community can supply the necessary common goods. BQ dynamics often describe a cheater-cooperator scenario, where some ecotypes, the “cheaters,” produce no common goods and rely on others, the “cooperators”, for survival. We have previously proposed that in systems with multiple common goods, an alternative endpoint for BQ dynamics can emerge. This endpoint describes an ecosystem of interdependent ecotypes engaging in “mutual cheating”, i.e. where common good production is *distributed*. However, even with multiple goods the common good production can be *centralized*, i.e. with one ecotype providing all common goods for the ecosystem. Here, we present an eco-evolutionary model that reveals that BQ dynamics can result in both distributed- or centralized common good production. The interaction range, *i*.*e*. the number of beneficiaries a producer can support, distinguishes between these two endpoints. While many ecosystems evolve to be *maximally distributed* or *maximally centralized*, we also find intermediate ecosystems, where ecotypes that appear redundant are coexisting for long periods of time. Due to the limited interaction range, these redundant ecotypes are unable to distribute the production of common goods fully due to the presence of non-producing types. Despite non-producers thus stalling the division of labor, we observe that sudden structural shifts can occur that purge the non-producers from the ecosystem. Overall, our findings broaden the understanding of BQ dynamics, unveiling complex interactions beyond the simple cheater-cooperator narrative.

## Introduction

Microbes often exhibit interdependencies to establish a community (Bordenstein and Theis, 2015; Cordero and Polz, 2014; Doolittle and Inkpen, 2018; Morris et al., 2013). These microbial interdependencies are especially likely to prevail in complex communities, such as soil and compost, where most ecotypes are unculturable (Albertsen et al., 2013; D’Onofrio et al., 2010; Iverson et al., 2012; Kim et al., 2019; Schwank et al., 2019; Starnawski et al., 2017; Stewart, 2012; Villanueva et al., 2017). In an interdependent community, interactions between microbes are often mediated through the production of common goods (also called public goods), such as secreted metabolites in cross feeding (Bruger and Waters, 2016; D’Souza et al., 2014; Driscoll and Pepper, 2010; Germerodt et al., 2016; Meijer et al., 2020; Morris et al., 2013, 2014). These microbial interactions build and maintain chemical cycles (Bonnefoy and Holmes, 2012; Doolittle and Inkpen, 2018; Morris et al., 2013) and regulate complex symbioses with larger organisms (Barko et al., 2018; Compant et al., 2019), including those with direct health impacts to human well-being (Kent et al., 2020; Partridge et al., 2018). Such interactions also have important consequences for the stability of microbial communities (Datta et al., 2016; Pande et al., 2014, 2016; Tekwa et al., 2017). Hence, understanding the key drivers underpinning microbial interdependencies will be important to predict the future of Earth’s complex ecosystems (Wortel, 2023; Wortel et al., 2018)

A growing framework for understanding how microbial communities evolve interdependencies is the Black Queen (BQ) hypothesis (Morris et al., 2012). The BQ hypothesis extends well-established socio-biological ideas, such as the ‘free-rider’ and ‘tragedy of the commons,’ to explain how microbes evolve interdependencies (Ndhlovu et al., 2021). The hypothesis posits that a certain function in a microbial system is a ‘leaky’ common good (Morris, 2015a; Morris et al., 2012), meaning that a beneficial function is provided that cannot be sequestered by the provider of the function. Examples of common goods are secreted compounds, such as many antibiotic resistance genes (*e*.*g*., beta-lactamases (Cairns and Jalasvuori, 2018)), elastases (Teufel and Gotz, 1993), and amylases (Baysal et al., 2003). Moreover, intra-cellular functions can apply. For example, Morris *et al*. report on a peroxidase that de-toxifies the surrounding milieu (Morris et al., 2012), and a similar effect is observed for heavy metal de-toxification (Boyd and Barkay, 2012; Parnell et al., 2010). Irrespective of the exact nature of the common good, when a common-good producer is surrounded by other producers, it will gain a selective advantage by ceasing production commensurate with the cost of production. When this common good is essential, a lack of common-good producers may result in community collapse. Thus, the BQ hypothesis predicts that there will be producers (‘cooperator’) trapped in providing everyone else (‘cheater’) with the common good.

The above cheater-cooperator prediction is sensible for systems with a single common good (Cairns and Jalasvuori, 2018; Cordero et al., 2012; D’Souza et al., 2014; de Mazancourt and Schwartz, 2010; Estrela et al., 2016; Germerodt et al., 2016; Lee et al., 2012; Morris et al., 2012; Ndhlovu et al., 2021; Pande et al., 2016, 2014; Shou et al., 2007; Stump et al., 2018; Van Vliet et al., 2022; Wang et al., 2019), and the prediction has been successfully applied to advance our understanding of microbial ecology and evolution (Brunet and Doolittle, 2018; Cairns and Jalasvuori, 2018; D’Souza et al., 2014; Derilus et al., 2020; Hesse and O’brien, 2024; Mas et al., 2016; Morris, 2015b; Morris et al., 2014; Ndhlovu et al., 2021; Takeuchi et al., 2024; Yuan and Meng, 2020). However, most communities in nature are likely to contain more than one common good (Morris, 2015a). In such communities, the BQ hypothesis permits other interdependencies to evolve. Indeed, a model assuming two common goods shows that a system can evolve, not only a cheater-cooperator pair as in a single-good system, but also a pair of producers, each producing a single good, depending on neighbour certainty and interaction range (Stump et al., 2018). An even wider variety of interdependencies become possible if, as is likely in nature, the number of common goods substantially exceeds one (**Figure 1A**). One end of this spectrum is the simple cheater-cooperator pattern, where one member retains all genes encoding leaky functions (the so-called “shooting-the-moon” strategy), and the other loses all these functions – the situation we hereafter call the *maximally-centralized* Black Queen (**Figure 1B**, left-hand side) (Morris, 2015a). The other end of the spectrum is the most extreme division of labor, where every member performs a single leaky function, and everyone relies on everyone else – the situation we hereafter call the *maximally-distributed* Black Queen (**Figure 1B**, right-hand side) (Fullmer et al., 2015). Between these extremes lie numerous other combinatorial possibilities. What interdependencies will a microbial ecosystem reach under what conditions?

**Figure 1.**
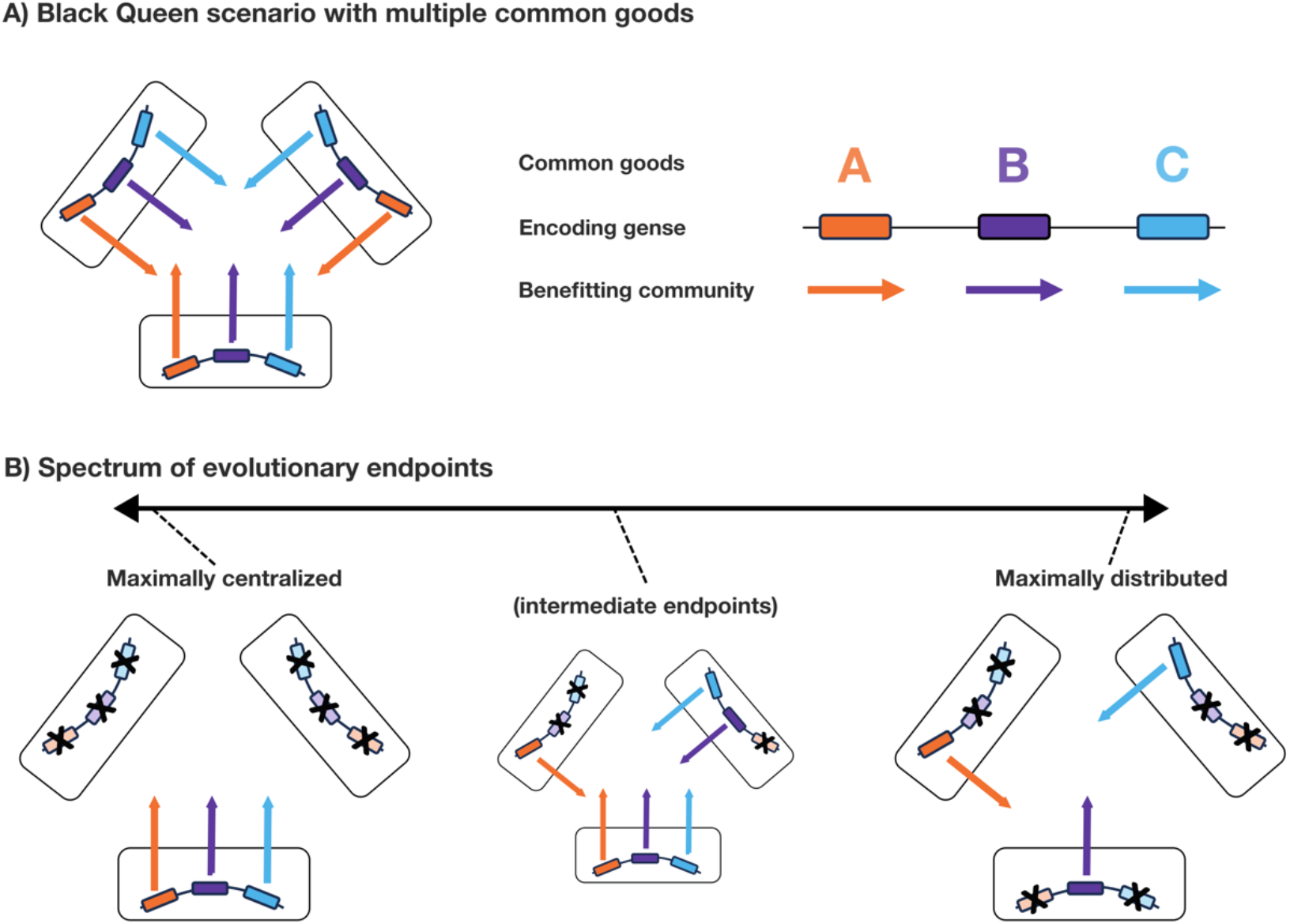
Schematic of Black Queen dynamics with multiple common goods and the spectrum of evolutionary endpoints. With multiple common goods, distinct evolutionary endpoints can be envisioned when starting with an initial population of omni-producers (shown in A). At the extremes of this spectrum of possibilities, the system can end up in one omni-producer with multiple cheaters (the *maximally centralized* Black Queen endpoint), or in a system where all members are trapped producing a minimal number of common goods (the *maximally distributed* Black Queen endpoint).

Here, we study an evolutionary model with many common goods and investigate the evolution of centralized-, distributed-, or intermediate BQ ecosystems. In our spatially explicit individual-based-model, we study ecosystems that emerge from a starting population of “omni-producers”, i.e., cells producing all the common goods. We identified two parameters that shape the interdependencies that evolve: i) the interaction range (i.e., the number of beneficiaries within the neighborhood of a producer) and ii) the cost to produce a common good. Depending on these parameters, both centralized- and distributed BQ ecosystems can emerge. Surprisingly, we found large areas of parameter space where common good production evolved to be only partially distributed. We show that this is due to the limited number of ecotypes that can fit within range of a certain producer, making complete division of labor statistically unfavorable. Finally, we show that the early emergence of non-producers (i.e. cheaters) can stall the further division of labor, simply by taking up space within the neighborhood. This results in ecosystems that are seemingly unchanging for millions of timesteps, which can nevertheless suddenly change in ecosystem structure. Taken together, our model shows how interaction range – and the resulting degree of neighborhood certainty – has a strong impact on microbial ecosystems driven by Black Queen dynamics.

## Methods

### Model overview

Our model is a two-dimensional toroidal grid of 200×200 grid points (**Figure 2A)**. Each grid point may hold a single individual (hereafter, cell). Each cell carries a genome of P loci that determines whether a distinct common good (CG) is produced or not. Each locus has an independent chance of deletion/inactivation of the gene (the difference between deletion and inactivation in cost to the host is negligible, so they are not differentiated) during every timestep. Each timestep, each cell rolls against the background death rate, and those that fail are removed, ensuring space for the survivors to grow into on the next timestep. Mixing of cells occurs through the Margolis diffusion algorithm (Toffoli and Margolus, 1987). During every growth phase of a timestep, open spaces will be competed for by neighboring cells (**Figure 2A / Figure S1)**. A cell must have access to all the CGs to reproduce, which can be achieved by self-producing the CGs or by being in the proximity of producers. If all CGs are available, a cell will reproduce with a baseline chance that is linearly decreased for each CG this cell produces. Thus, cells that produce fewer CGs will successfully reproduce more often, provided that they have access to all CGs. We do not explicitly model the CGs themselves, and no diffusion and decay of CGs is considered in our model. Instead, these processes are assumed to be in steady state, with access to CGs only being determined by the neighborhood size.

**Figure 2.**
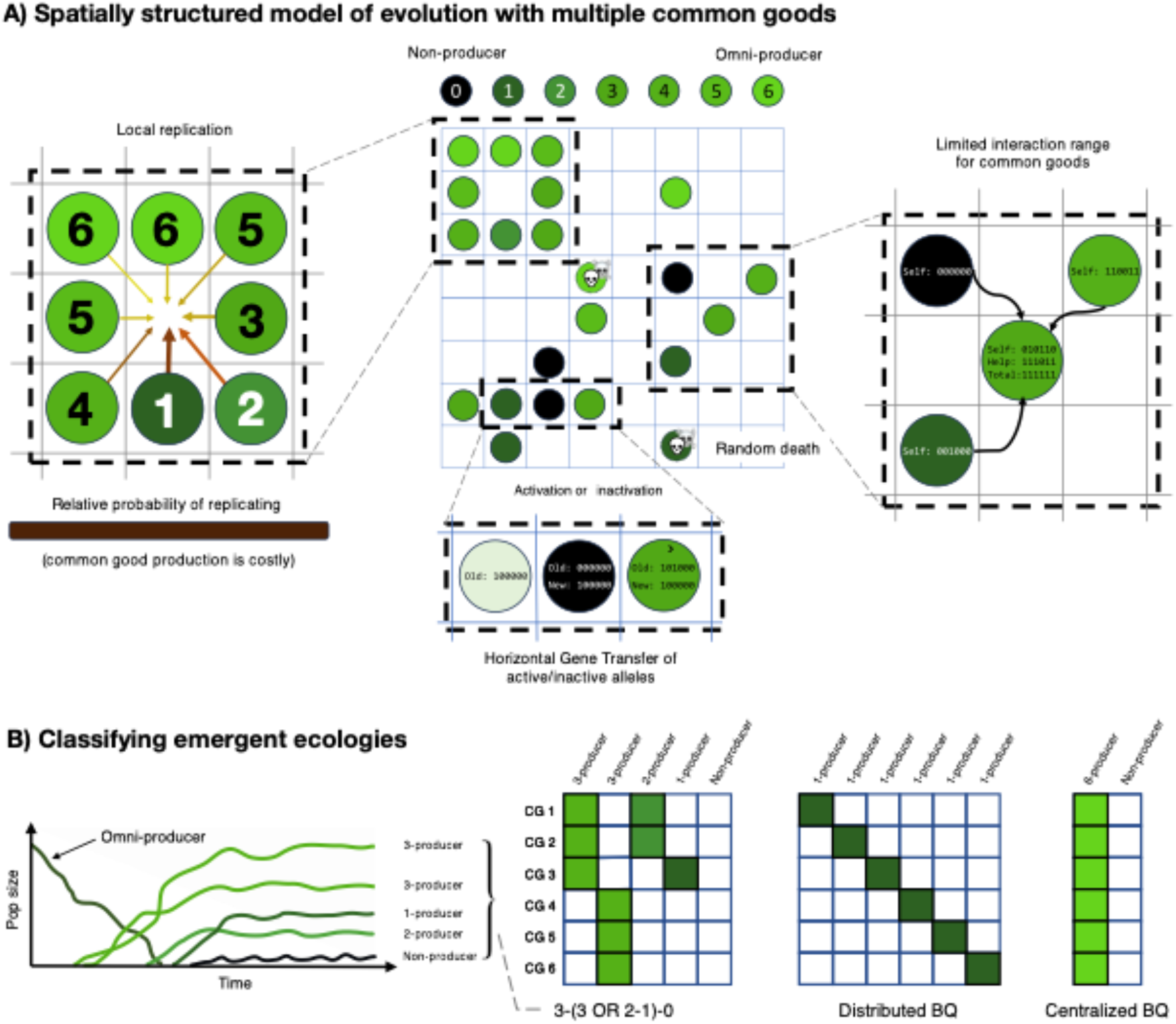
**A) Spatially structured model of evolution with multiple common goods (CGs).** All cells in a Moore neighborhood of an empty space will have a chance to reproduce into that space. Cells that produce fewer goods have a greater chance of reproduction, provided they have access to all CGs. Cells within a certain interaction range can provide CGs to the focal (center) cell. HGT can change the genomic loci of neighbouring cells. This includes gaining new genes (activation), but also functional alleles being overwritten by non-functional alleles (inactivation). The model is implemented in Cacatoo, with an interactive version is available online (https://mfullmer.github.io/). The source code is available on github (https://github.com/MFullmer/BlackQueen). **B) Classification of emergent ecologies**. An illustrative graph showing how an initial omni-producing type can evolve into a complex 3-(3 OR 2-1)-0 ecosystem. The adjacent table shows how a 3-producer could pair with the either the other 3-producer or with the 2- and 1-producer to gain access to all 6 CGs. The other two tables illustrate maximally distributed- and maximally centralized BQ endpoints.

### Software

Simulations were written in JavaScript using the module *Cacatoo* (van Dijk, 2022). Data visualization used the R (R Core Team, 2020) package ggplot2 (Wickham, 2016). An interactive version is available online (https://mfullmer.github.io/). The source code is available on github (https://github.com/MFullmer/BlackQueen).

### Horizontal Gene Transfer

The impact of Horizontal Gene Transfer (HGT) may significantly impact the evolution of interdependencies, as it enables non-producing ecotypes to recover the production of one or more CGs. We therefore included HGT in our model as a chance to overwrite an allele at a certain locus with the corresponding allele from a neighboring cell (**Figure 2A**). This means that genes can be activated/gained through HGT, but can also be inactivated through HGT. Genes/alleles are not linked and each one transfers independently of the others.

### Ecotype nomenclature

We choose to use the term “ecotype” to denote a specific “strain” of microbe in an ecosystem over the potentially synonymous “genotype.” Although genotype is also appropriate in our model, which incorporates the specific genetic background that creates the ecologically-relevant phenotype, we made this choice because we are interested in the ecosystem and interdependencies over the genetic details. We denote an ecotype producing *m* kinds of CGs as “*m*-producer.” We refer to an ecotype producing all the system’s common goods as an ‘omni-producer’ (i.e., *n*-producer in a system requiring *n* CGs for growth). We refer to an ecotype that produces no CG as a ‘non-producer’ (or 0-producer). Although non-producers could be referred to as “cheater”, one could also say that a 2-producer is a cheater *relative to* a 3-producer. To avoid this ambiguity, we avoid the word “cheater”.

### Ecosystem nomenclature

In our model, complex microbial ecosystems can evolve. We describe the states of evolved ecosystems as numeric strings representing the division of labor between ecotypes (**Figure 2B**). For example, 3-3 describes an ecosystem comprised of two 3-producers equally dividing production of 6 CGs. In a similar fashion, 4-2 describes a system with an asymmetric division of labor. If a third ecotype exists that produces no common goods, the ecosystem string would be 4-2-0. When an ecotype produces common goods that are also provided by other ecotypes, we put the numbers between brackets with logical operators. For example, 3-(3 OR 2-1) describes a community where the left 3-producer could pair with either the 2-1 pair or the other 3-producer to access the 6 CGs. For clarity, we provide table-representations to show how CG production is divided amongst ecotypes, which is especially helpful when the ecosystem cannot uniquely be specified by the string notation.

### Model initialization and parameters

We initialize our ecosystem as a population of omni-producer cells. Then, we run long-term evolutionary experiments for at least six million timesteps (approximately 600,000 generations) for the main results, and two million timesteps in our simulations with HGT. An overview of the model parameters and their values is shown in **Supplementary Table 1**.

### Competitions & Invasions

Competition and invasion experiments were carried out with mutation and horizontal gene transfer disabled. Two separate grids are initialized, one with one pre-selected competitor combination of ecotypes and the other with the second competitor combination. For example, one grid might have a 3-3-0 ecosystem while the other has 3-2-1. Each grid was allowed to equilibrate for several thousand timesteps, typically 10,000, in isolation. After this equilibration we initiated the invasion through the methods explained in the next paragraph. After the invasion step, typical simulations then ran another 80,000 steps to reach 100,000 total. Some experiments were run longer to ensure the outcomes were stable in the long-term.

In our method of invasion, a region of one grid is copied onto the corresponding region of the second grid, and vice versa. This invasion by region ensures that the invader ecotype(s) start in proximity to any essential partners. The area of grid transferred was typically 1/16^th^ of the total system. However, for some experiments we used up to a 50:50 mix. We also tested a second method of invasion that swaps random points on the grids. This reduces the likelihood that the invading ecotypes will have proximity to essential partners from their own grid, but they may have access via the occupants of the new grid. This invasion-by-point method yielded largely similar results to invasion-by-region, so its data is not presented.

We also implemented two types of special invasion experiments to gain specific insights. The first uses “stalling blocks” to simulate the space-filling effects of non-producers. These were spawned into the simulation concurrently with the invasion from the parallel grid. These blocks occupy space on the grid but they do not reproduce or die. They also produce zero common goods. These are spawned in at random points to prevent unrealistic effects from regional replacement. Our second special invasion turns the 6a ecotype in the 6-(6 OR 4-2) ecosystem into a “stalling block-like” actors. These 6a are allowed to produce common goods. However, it was only subject to death when its proportion of the community rose above the set value. Likewise, it was only allowed to grow when below. This allowed it to behave almost identically to a “normal” 6N while the overall proportion remains essentially fixed over time. The other three ecotypes were initialized to their “natural” proportions relative to each other and allowed to equilibrate to their new balance points. By varying the starting proportions of this block-like 6N, we could examine the frequency-dependence of 6R and 4-2.

### Phase Diagrams

To show the division of CG production as a function of the interaction range (area within which CGs are shared) and cost per CG, we made phase diagrams (**Figure 3 and Figure 4**). In these figures, the first line of text in each square shows the state of an ecosystem consisting of the most dominant set of ecotypes that together supply all CGs. This was determined by identifying the most numerous producers of each CG and then summing the total production of all those ecotypes. The 2^nd^ line of text shows the next most common alternate set of ecotypes that together supply all CGs. Only CGs with a 2^nd^ most common producer that supplies at least 25% of at least one CG’s total production are counted. “None” is written if there are no alternate set of ecotypes supplying at least 25% of at least one CG. The color of each block shows the proportion of all CG production accounted for by the combination printed in the first line. Yellow colors indicate that there are very few ecotypes not accounted for, while purples indicate that the printed combination is only a small fraction of the total population.

**Figure 3.**
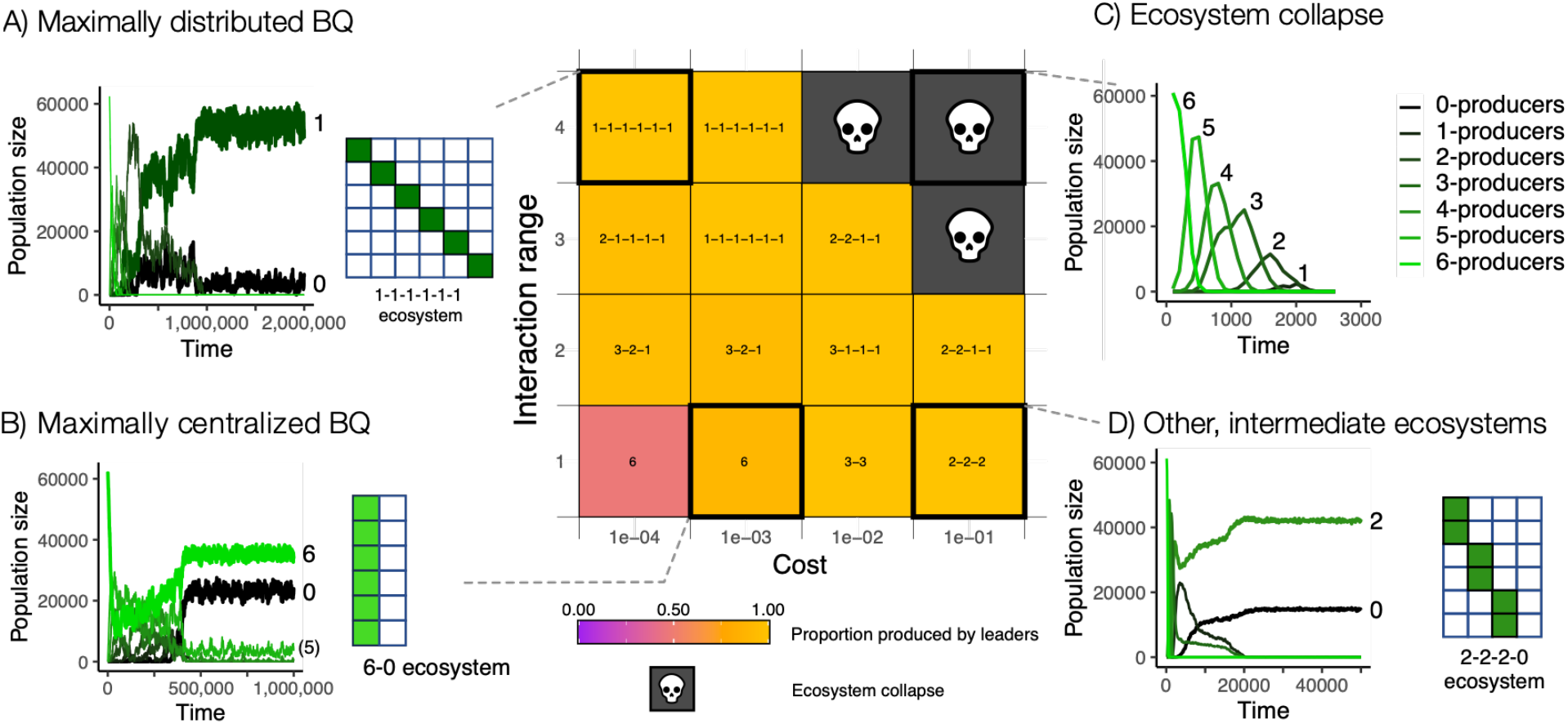
Phase diagram showing results for combinations of interaction range and production cost with 6 common goods (CGs). There is no HGT in these simulations. We first determined which ecotypes produces the largest amount of each CG (called “leaders”), and the total production of these leading ecotypes. The text in each tile shows this primary combination. **A) Exemplar of maximally distributed BQ endpoint** observed when interaction range is large and CG production costs are low. **B) Exemplar of maximally centralized BQ endpoint** when interaction range is small, and costs are low. **C) Exemplar of dynamics showing population crash** promoted by the high costs of CG production, resulting in the “tragedy of the commons”. **D) Exemplar dynamics showing intermediate ecosystems**, neither maximally distributed or centralized.

**Figure 4.**
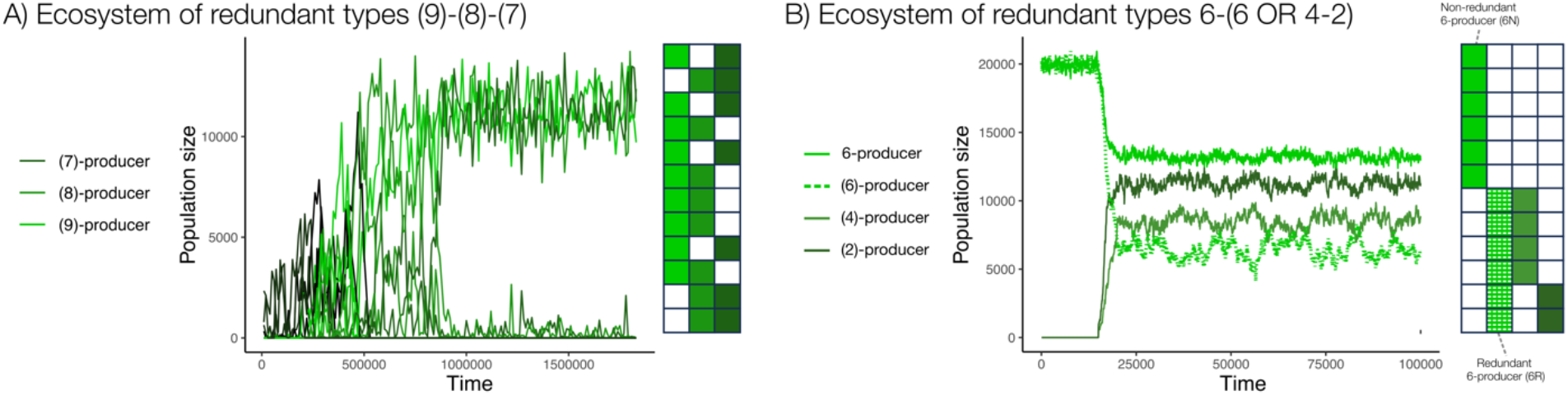
Examples of the evolution of redundant ecotypes in simulations with 12 CGs. Redundant ecosystems evolve when the cost of CG production is high and the interaction range is low (shown for 6 CGs in Figure 3D). **A) Example of complete redundancy**, where any pair of the three ecotypes can complete a full suite of CGs. **B) Example of partial redundancy**, where one non-redundant ecotype is responsible for the production of the first 6 CGs, whereas the production of the remaining 6 CGs is divided among three redundant ecotypes.

### Data availability

Simulation data from figures and supplementary figures is available online at (https://github.com/MFullmer/BlackQueen_Data).

## Results

### Interaction range distinguishes between two modes of Black Queen dynamics

To understand what interdependencies evolve when microbes need many common goods (CGs) to grow, we first investigated our model by sweeping across parameters (see Methods). Two parameters emerged as key determinants of the model’s outcome: the cost of CG production and the interaction range (see **Figure 3**). Simulating with 6 CGs, we found that when the costs of CG production are low, both distributed- and centralized BQ ecosystems can evolve, depending on the interaction range. When the interaction range is large, ecosystems evolve to be maximally distributed (**Figure 3A**), with each ecotype producing a single CG (this ecosystem is denoted as a 1-1-1-1-1-1). When interaction range is small, ecosystems evolve towards a maximally centralized solution (**Figure 3B**), with a single lineage producing all 6 CGs (hereafter, an omni-producer) accompanied by a non-producer (this ecosystem is denoted as 6-0). The latter ecosystem fits the classic cheater-cooperator framework. However, when interaction range is intermediate or the costs of CG production are high, ecosystems evolve to be neither maximally distributed nor maximally centralized. Instead, ecosystems display partial division of labor, with one or more ecotypes producing multiple CGs (**Figure 3**; e.g., 3-3-0 ecosystem when the cost is 10^−2^ and the interaction range is 1). Moreover, in many cases, ecosystems display a surprising degree of functional redundancy. For example, the 2-2-(2 AND 1) ecosystem shown in **Figure 3D** illustrates that one of the 2-producers overlaps with a 1-producer in CG production. This raises the question as to why the 2-producer does not give up production of the redundant CG. When simulating with 12 CGs, we found an ecosystem where each of the three ecotypes (namely, 9-, 8-, and 7-producers) could pair with one of the other two (**Figure 4A**). Even more surprisingly, we also observed a 6-(6 OR 4-2) ecosystem, where the 4- and 2-producers together produce the exact same CGs as the 6-producer (**Figure 4B**). Thus, the other 6-producer can pair up with either the 6-producer, or the 4-2 pair. Taken together, these results indicate that microbial ecosystems can evolve a wide variety of interdependencies and functional redundancy depending on the interaction range and the CG production cost.

### HGT has minor effects on ecosystem structure

HGT enables microbes to recover the production of one or more CGs, so it may significantly impact the dynamics in our model. To investigate this effect, we repeated the above parameter sweep with the addition of varying rates of HGT (see methods). Interestingly, while we observed a strong effect on diversity and therewith the CG production by the dominant ecotype, the evolved ecosystems are remarkably similar (**Figure 5**). Even when HGT outpaces the mutation rate 100-fold, distributed and centralized BQ ecosystems evolve in the same regions of parameter space. That said, there are some differences. At high rates of HGT there are many more uncommon ecotypes. This results in the leading producer types tending to comprise a much smaller portion of the total population (purple instead of yellow squares in Figure 5). As a result, some ecosystems now have more multi-producers present (e.g., the ‘2s’ in a 2-2-1-1-1-1 system) among the leaders than without HGT. However, we found that at any given time, these multi-producing cells are the primary producer of only one CG. This means that the production of a second CG is likely incidental to a recent HGT event rather than a selectively advantageous trait. Taken together, the continuum of distributed- and centralized ecosystems is largely robust to the effects of HGT.

**Figure 5.**
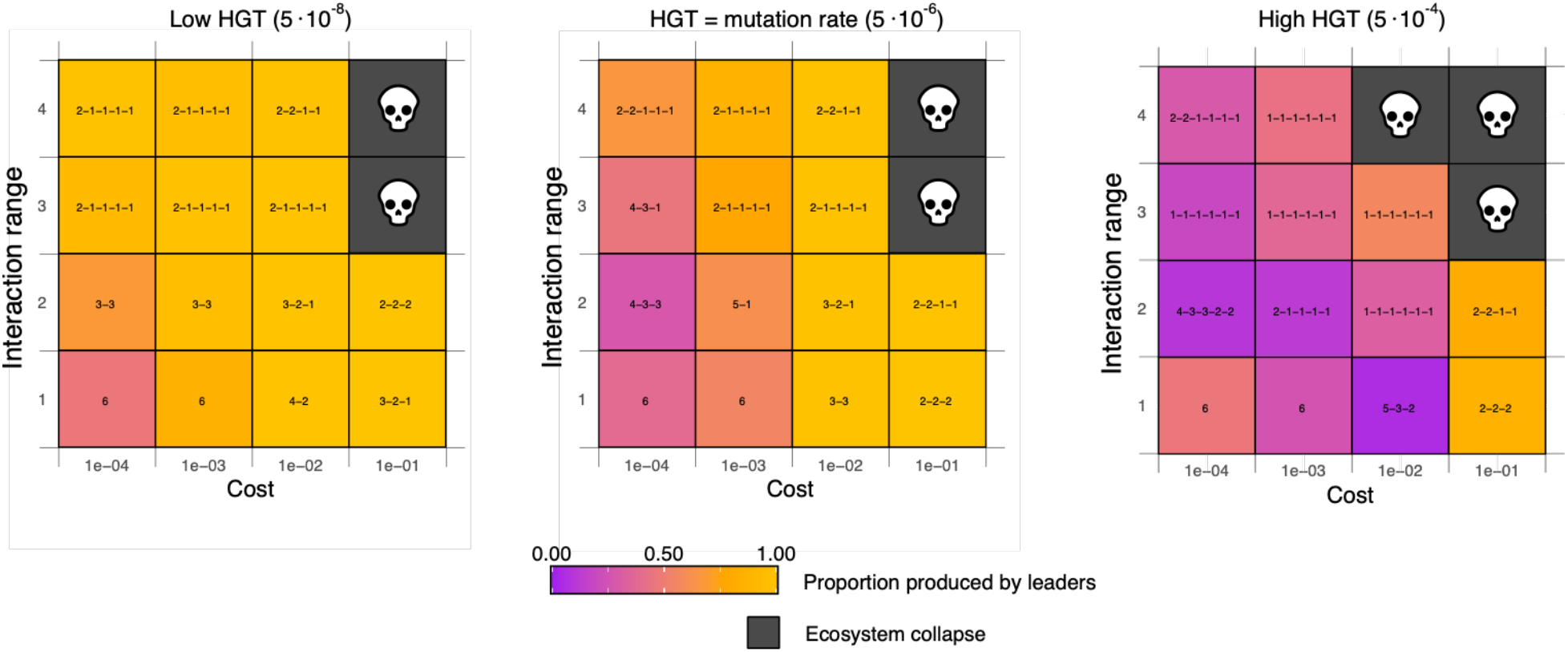
Phase diagrams with 6 common goods repeating the experiments shown in Figure 3 with HGT enabled, with low rates of HGT (5e-08, 100-fold lower than the mutation rate), HGT equal to the mutation rate (5e-06), and high HGT rates (5e-04, 100-fold higher than the mutation rate). Like in Figure 3, the top line shows the combination of types producing the largest amount of each CG (called “leaders). The background color indicates the proportion of CGs produced by the leaders.

### Redundant ecotypes fill distinct statistical niches

Above, we have discussed that high costs and limited interaction range often resulted in the evolution of redundant ecosystems, such as the 6-(6 OR 4-2) ecosystem. The redundant 6-producer and the 4-2 pair would seem to be, at face value, direct competitors, filling the exact same niche in the ecosystem. This raises the question as to why there is no competitive exclusion for millions of time steps, suggesting either a fully stable or at least quasi-stable system. To understand the coexistence of the seemingly redundant ecotypes, we performed invasion experiments. We invaded the redundant 6-producer (hereafter: 6R) into an ecosystem of the non-redundant 6-producer (hereafter 6N) and the 4-2 pair. We also performed the inverse and invaded the 4-2 pair into an ecosystem of the two 6-producers (**Figure S2**). Interestingly, we found that 6R and the 4-pair could both invade one another (i.e., the redundant 6-producer was able to invade the 6-4-2 ecosystem, and the 4-2 pair was able to invade the 6-6 ecosystem). These results indicate that 6R and the 4-2 pair are engaged in negative frequency-dependent dynamics that prevent competitive exclusion. These dynamics might arise from two commensal relationships existing in parallel: either a 4- or 2-producer free-rides on the CGs produced by the 6R-producer. Each of these relationships is essentially a simple cooperator-cheater relationship, but they are embedded in an intricate web of interdependencies, which, as a whole, could ensure the coexistence of all ecotypes (**Figure S3**).

We hypothesized that the apparent 6R and 4-2 frequency dependence was mediated by a combination of selective advantage through reduced costs and likelihood of residing near a complementary producer. To test this, we created simulations of the (6-(6 OR 4-2)) system where the frequency of 6N (the 6-producer that can complement with either 6R or the (4-2) pair) is fixed. Comparing the proportions of 6R to 4 and 2, we find that (4-2) relatively prospers when 6N is rare and that 6R does so when 6N is most abundant (**Figure S4**). This result reinforces the conclusion that 6R and 4-2 are negatively frequency dependent and helps explain how the system is stable, or at least quasi-stable. As the cost of CG production is steep (10% chance of reproduction penalty per CG) it would be expected that ecotypes that produce fewer would be advantaged. This is observed at low frequencies of 6N. However, when 6N is highly numerous and begins to physically occupy much of the space in the simulation, 4 and 2 begin to dwindle; and even go extinct at the extremes. We can understand this result by considering that because of the small neighborhood size, the 6-producer gains a statistical advantage; it is more likely to be adjacent to the 6-producer than the (4-2) pair are to be next to each other *and* the 6-producer. Our results show that these two opposing factors, i.e. the selective advantage of not producing a CG and the statistical likelihood of being next to a producer, can balance each other out.

### Non-producers stall the division of labor

In the previous paragraphs, we described how complex ecosystems with redundant ecotypes could emerge and remain in “quasi-stable” states for indefinite periods while remaining recalcitrant to invasion by ecotypes that would seemingly be advantaged (i.e., non-producers being unable to invade the (6-(6 OR 4-2)) system). However, these redundant systems were not the only examples of invasion-resistance observed. We found multiple cases of evolved ecosystems spending millions of timesteps in a single state before suddenly undergoing a major change to the ecosystem composition. In one example of note, a 3-3-0 system arose and persisted for ∼4 million timesteps before suddenly transitioning into a 3-2-1 ecosystem (**Figure 6**). Notably, the non-producer (0-producer) ecotype is very rare after this transition, constituting no more than a few percentage points of the total. This reduction in non-producers is likely due to the above-mentioned statistical effects of being next to producers, which is significantly reduced in the 3-2-1 ecosystem, hence disfavoring non-producers. But why did it take such a long time for the ecosystem to transition into the 3-2-1 ecosystem?

**Figure 6.**
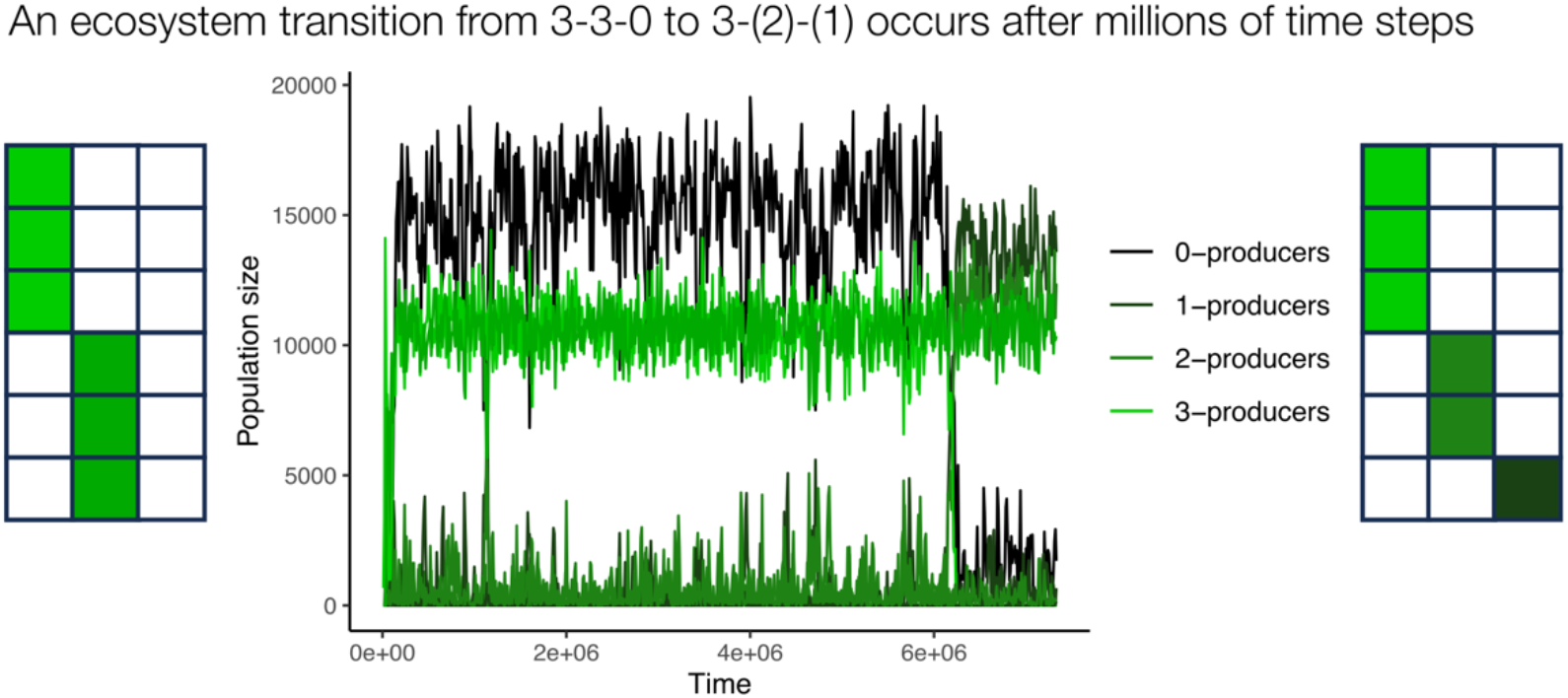
Black Queen dynamics can be halted for millions of time steps. A 3-3-0 ecosystem is seemingly stable for 6 million time steps, and then transitions to a 3-2-1 ecosystem, where the non-producers are all but pushed out.

We hypothesized that the difficulty to transition from a 3-3 to a 3-2-1 ecosystem could also be explained due to the statistical effects as described above. When non-producers are present, it may be more difficult for the ecotypes to divide labor due to the reduced likelihood of meeting. These effects are illustrated in (**Figure 7A-B**). In other words, while non-producers are themselves the result of BQ dynamics, they appear to have a stalling effect on the evolution of even more distributed Black Queen ecosystems.

**Figure 7.**
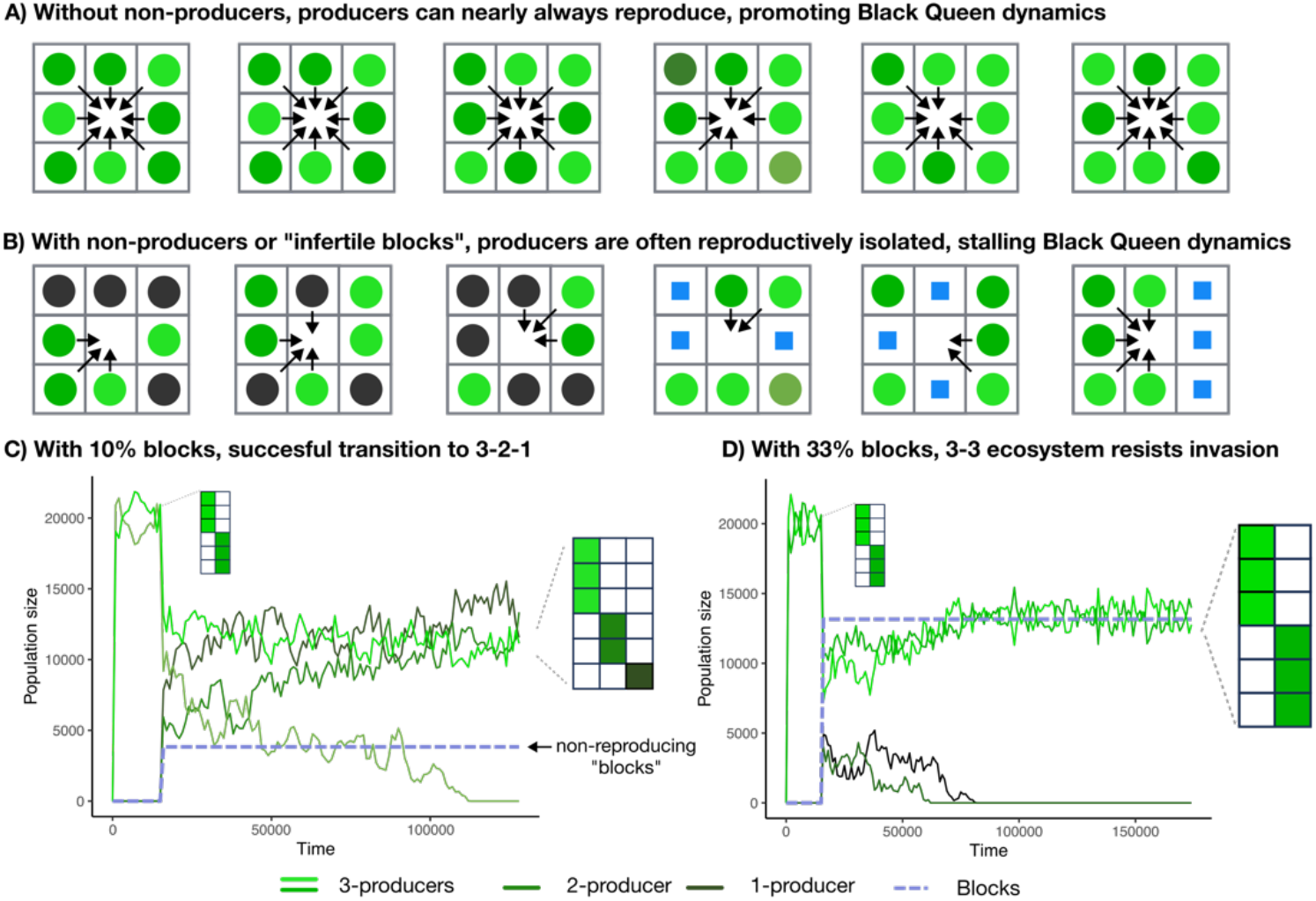
The presence of infertile “blocks” illustrates the effects of neighborhood certainty. **A)** In the presence of 10% infertile (not reproducing) blocks, a 3-3 pair is successfully displaced by a 3-2-1 set. **B)** With 33% blocks, the 3-3-0 ecosystem is not displaced by the invading 3-2-1 set.

To better understand the stalling effects of non-producers, we implemented a special deletion that converts any ecotype (regardless of how many CGs they produce) into a non-producer with a single mutation. These rare events allowed 0-producers to emerge much more rapidly appear from the initial population of omni-producers. We observed that timing played a key role in the resultant community.

When an early mutation caused the early emergence of non-producers, they indeed appeared to stall the further evolution towards more distributed ecosystems. In the most extreme cases, a very early emerging -producer would even freeze the system in a (12-0) ecosystem state, the classic cheater-cooperator pair, with no further change (**Figure S5A**). In contrast, when the non-producers arose late, for example after a redundant system (such as 6-(6 OR 4-2)) had already developed, they failed to establish and were eliminated from the system (**Figure S5B**). These simulations suggest that non-producers and partial producers are engaged in a positive frequency-dependent interaction that constrains future community evolution.

### Neighborhood certainty drives the “stalling” effect of non-producers

There are two candidate explanations for the stalling effect of non-producers. First, non-producers may have a selective advantage in reproduction, meaning they continuously outgrow the other ecotypes. When spaces open through death, the non-producers replicate into it much more readily than ecotypes burdened with larger shares of CC production. This slows or prevents the growth of the heavier-producing ecotypes, possibly allowing random death to purge them if they are rare. Alternatively, the stalling effect may be acted out through crowding effects. By simply occupying space on the grid, non-producers can reduce the neighborhood certainty for partial-producing ecotypes, a statistical effect described in detail above. This would then create a selective force towards ecotypes that can produce a greater variety of CGs to increase the statistical likelihood of forming complete sets of goods with one’s neighbors. To distinguish between these two explanations, we experimented with the 3-3-0 to 3-2-1 eco-evolutionary transition mentioned above. We dissected the mechanism by repeating the 3-3 to 3-2-1 invasion experiments in the absence of the non-producer, but in the presence of “stalling blocks” (hereafter: blocks), at the time of invasion. Blocks occupy space on the grid, but do not produce goods, reproduce, or die. Herewith, we eliminate the selective advantage of non-producers, and are left only with the fact that non-producers take up space. If blocks do not prevent 2-1 from invading in the same fashion as non-producers, that indicates that the reproductive advantage of the non-producers drives the stalling effect. Contrarily, if the 2-1 invasion fails, then this indicates that the spacial occupancy of the non-producers drives stalling. We found that the simple act of taking up space was sufficient to explain the stalling effect of non-producers. When the number of blocks is sufficiently high (33% of total grid spaces), 3-3 wins over 3-2-1 100% of the time (n=5) (**Figure 7C**). When the number of blocks were dropped to 10%, (3-2-1) wins the competition 100% of the time (n=6) (**Figure 7D**). This reveals that the non-producer’s reproductive advantage has only minor effects, and that the occupation of space is the most important variable of non-producers modifying the interaction structure in the ecosystem. In summary, statistical effects such as neighborhood certainty can be important drivers of the evolutionary trajectories in spatially structured ecosystems.

## Discussion

We have investigated simulations of digital bacteria-like organisms undergoing the evolution of interdependencies through Black Queen Dynamics. Our simulations show that two distinct outcomes of the Black Queen dynamics can be attained in systems featuring multiple common goods. In one outcome labor is divided between an “omni-producer” (producing all the common goods) and a non-producing cheat (which we call the *maximally centralized* Black Queen endpoint). In the other outcome, labor is divided between many lineages which all produce the minimal amount of common good (the *maximally distributed* Black Queen endpoint). Interestingly, these two outcomes are not determined by the cost of CG production, but are predominantly driven by the interaction range, a possibility that was coined in previous work using two common good systems (Stump et al., 2018). When interaction range is large, many ecotypes can fit within one neighborhood and completely divide labor (*i*.*e*. distributed BQ endpoints are favored). With a smaller interaction range, we observe centralized BQ endpoints that fit the classical cheater-cooperator narrative, as there is no room for additional dependencies within a single neighborhood. Interestingly, we found that large areas of intermediate parameter space resulted in neither of these two extreme BQ outcomes. Instead, we find complex ecosystems of partially or completely redundant ecotypes. While these redundant ecosystems are remarkably stable, sudden transitions are occasionally observed after hundreds of thousands of generations. We found that these transitions are governed by complex frequency-dependent exclusion mechanisms between non-producers and partial producers, mediated by the limited neighborhood sizes in our spatially explicit model. Thus, interaction range is an important driver of BQ dynamics, whether it be distributed, centralized, or otherwise.

Prokaryotes are known to evolve mostly through changes in gene content. Gene gain is primarily driven by the process of horizontal gene transfer (HGT). However, much theory of BQ dynamics does not yet include this mechanism. In our simulations, we investigated how ecosystems responded to rampant rates of HGT. We found that the ecosystem structures that evolve under various BQ hypotheses are surprisingly robust to HGT, even when this outpaces mutations 100-fold (**Figure 5C**). This suggests that the process of HGT does not hinder the development of microbial communities with many metabolic dependencies, which indeed appear ubiquitous in nature (Kost et al., 2023). Hence, with or without HGT, BQ dynamics are likely of great importance to understand the complexity of microbial ecosystems. However, as we have here only modelled HGT in the abstract, future work could reveal a more interesting interplay between HGT and Black Queen dynamics. For example, when taking mobile genetic elements into consideration, one may study how different genes have different rates of gene transfer, a process that earlier modelling has shown to have substantial impact on microbial population dynamics (Van Dijk et al., 2020; van Dijk and Hogeweg, 2016).

Our model treats all common goods as equal, with all providing the exact same benefit to all cells in the vicinity. This approach neglects the ways in which a producer derives more benefit from production than nearby cells (Driscoll and Pepper, 2010). This is especially important when considering that the range at which a common good benefits others, may vary (Van Vliet et al., 2022). A secreted metabolite will diffuse much faster in a mixing flask than it will in a stagnant biofilm, and local uptake can deny cells any benefit if they are more than a few microns away (Co et al., 2019). These differences in range are further shaped by the specific leakiness of the common goods, (e.g., a secreted versus an intracellular enzyme) (Morris, 2015b). Such differences in the interaction rate can have very surprising consequences, for example with short-ranged interactions resulting in more common good production when the costs are increased (Colizzi and Hogeweg, 2016). Logical extensions of our work will therefore incorporate these factors. However, by excluding these other factors, our simple model clearly illustrates the basic principles of division of labor and frequency-dependent exclusion in ecosystems governed by multiple common goods.

While BQ dynamics are often understood in terms of the costs of producing common goods (D’Souza et al., 2014; Fullmer et al., 2015; Morris et al., 2012), our model shows that even minor costs results in the rapid evolution of microbial interdependencies. This raises the interesting possibility of neutral evolution being a significant driver of community outcomes. Indeed, when gene loss outpaces gene gain as suggested by genomics studies (Puigbò et al., 2014), one could reasonably expect BQ dynamics to play out even when costs are insignificant. In a similar line of reasoning, we have shown in **Figure 7** that the reproductive advantage of “cheaters” – which may benefit them individually – is not the reason why they shape the ecosystem structure. Instead, the simple fact that they take up space is an important consideration, a component that most models do not consider. Indeed, the partial and complete redundancies observed in our work do not appear to adhere to the optimal divisions of labor we expect from selection. These examples evoke thoughts of constructive neutral evolutionary processes, such as *“It’s the Song Not The Singer*” (Doolittle and Booth, 2017; Doolittle and Inkpen, 2018). In other words, we observe the evolution of complexity not because of adaptive reasons on the level of individuals, but because of the differential persistence of ecosystems (Lenton et al., 2021). Taken together, our results indicate that through very simple interactions, biological complexity that goes beyond the simple cooperator-cheater framework, emerges free from individual selection.

## Notes

https://github.com/MFullmer/BlackQueen_Data

https://github.com/MFullmer/BlackQueen

https://mfullmer.github.io/

### Competing Interest Statement

The authors have declared no competing interest.

