## Supplemental Figures and Tables for "Interaction range of common goods shapes Black Queen dynamics beyond the cheater-cooperator narrative"

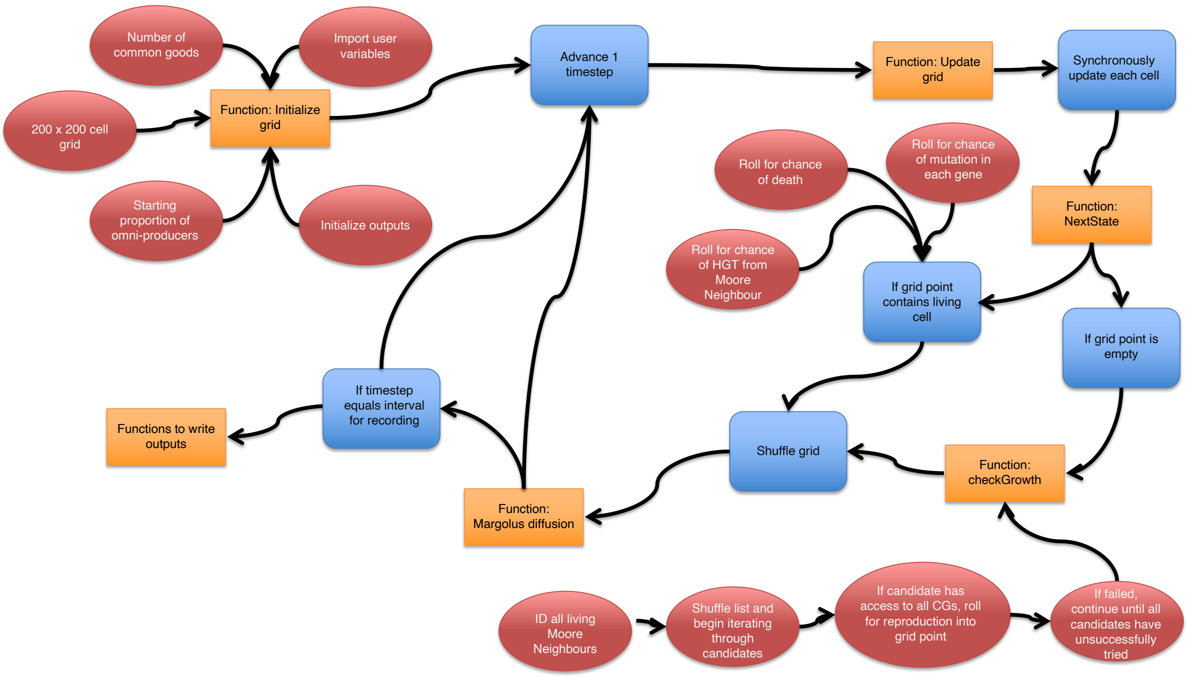


**Figure S1.** Workflow of the main simulation model.

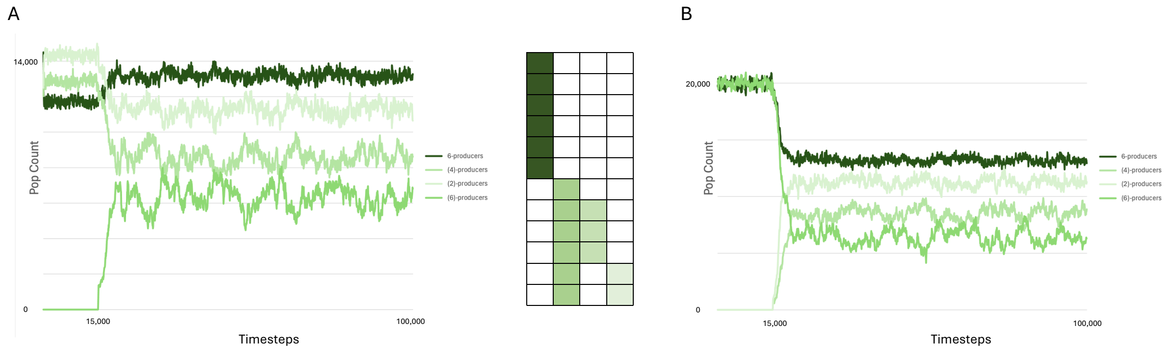


**Figure S2.** Invasions simulation results with the 6-(6 OR 4-2) system. Simulations were setup with paired grids, one with a 6-6 and another with 6-4-2 at the proportions observed in the "wildtype" equilibrium. Mutation was disabled. After 15,000 timesteps of equilibration the invasion was triggered by swapping parts of the two grids. **A)** 6 into 6-4-2. **B)** 4-2 into 6-6.

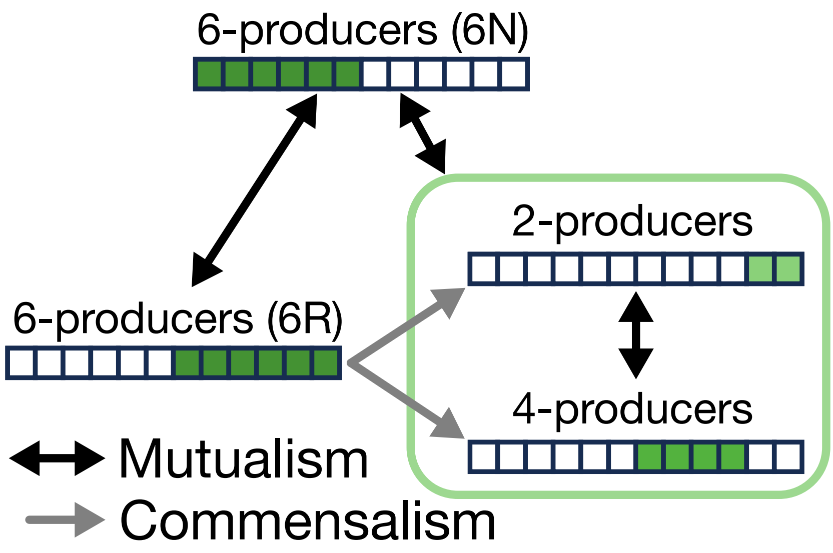


**Figure S3.** Interactions between the ecotypes in the 6-(6 OR 4-2) system.

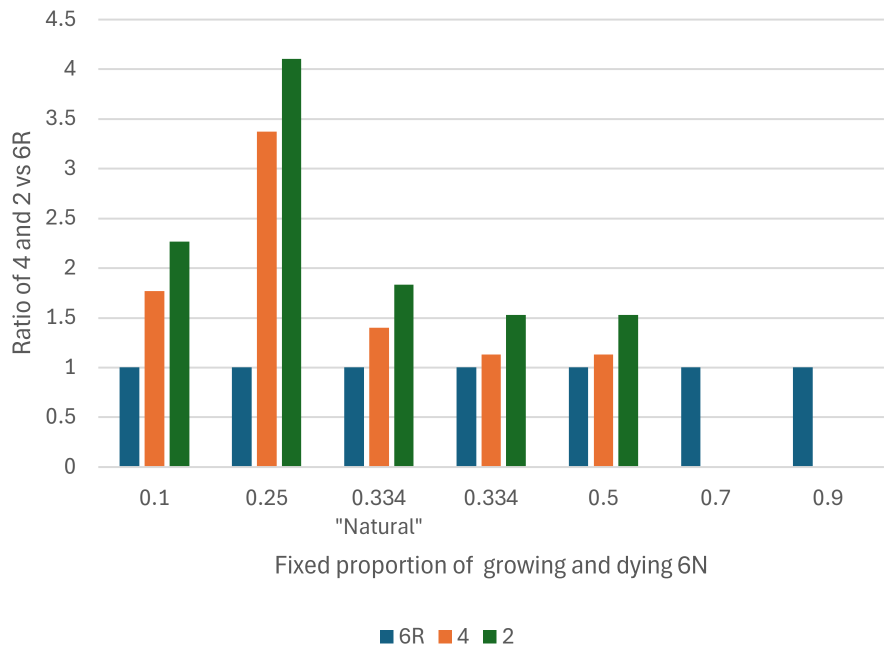


**Figure S4.** 6N as living blocks. A modification of the 6-(6 OR 4-2) system. 6N is altered to only grow or die when below or above, respectively, to maintain a fixed proportion of 6N cells. 6R, 4, and 2 are initialised at their equilibrium proportions to each other using the amount of free space after initialising 6N. Bars show the ratios of 4 and 2 against 6R, to allow for changing absolute numbers with different numbers of 6N.

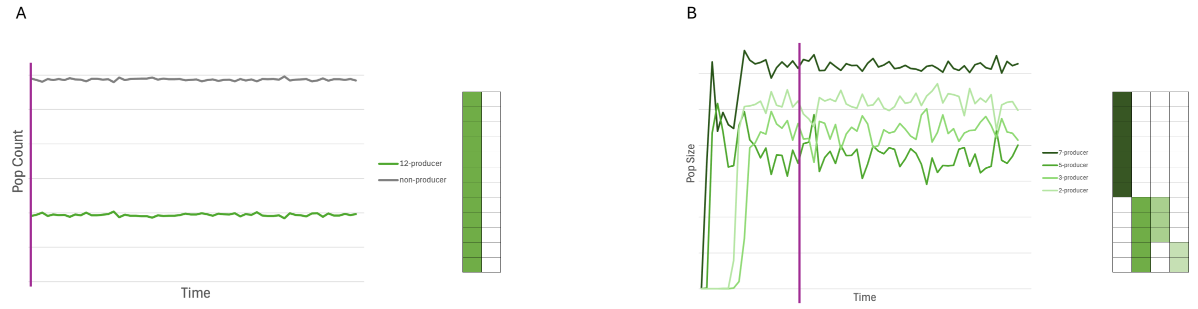


**Figure S5.** Simulations with super deletions. Simulations started with 12 common goods, a CG cost of 1E-02, default mutation rate and no HGT. A special super deletion parameter was added which allows all production to be converted to non-production in a single event (5.00E-09), i.e., the cell becomes a pure non-producer. **A)** Example of an early superdeletion event. Non-producers rapidly overtake the system, trapping it in a 12-0 configuration. **B)** Example of a late superdeletion event. The system had already formed a quasi-stable 7-(5 OR 3-2) configuration and the non-producer was unable to invade.

| **Default Parameters** | **Default values** |
|---|---|
| Grid Size | 200x200 cells (40,000 carrying capacity) |
| Baseline reproduction chance | 0.6 |
| Baseline death rate per timestep | 0.1 |
| Number of common goods (CGs) | 6 or 12 |
| Growth penalty per CG produced (Cost per CG * Baseline reproduction chance ) | 1e-04, 1e-03, 1e-02, 1e-01 |
| Instantaneous chance of reproduction | Baseline reproduction chance (0.6) - (#CGs produced * Growth penalty per CG produced) |
| Interaction range (Moore neighborhood radius) | 1 / 2 / 3 / 4 (radius of moore neighbourhood) ; 9 / 25 / 49 / 81 (number of potential beneficiaries, producer included) |
| Mutation rate / per gene / per timestep | 5e-06 |
| Horizontal Gene Transfer rate | 0 |

**Table S1.** Default model parameters.
